# Cross-sectional analysis of the microbiota of human gut and their direct environment (exposome) in a household cohort in northern Vietnam

**DOI:** 10.1101/2021.02.23.432416

**Authors:** Vu Thi Ngoc Bich, Ho Bich Hai, Gianluca Galazzo, Vu Tien Viet Dung, Melissa Oomen, Trang Nghiem Nguyen Minh, Tran Huy Hoang, H. Rogier van Doorn, Heiman FL Wertheim, John Penders

## Abstract

Comprehensive insight into the human gut microbiota and the interaction with their environment in communities with a high background of antibiotic use and antibiotic resistance genes is currently largely lacking. In a cohort (Vietnam), individuals within the same household, also individuals within their geographical cluster share more bacterial taxa than individuals from different households or geographical clusters. The microbial diversity among individuals who used antibiotics in the past four months was significantly lower than those who did not. Fecal microbiota of humans was more diverse than non-human samples, shared a small part of its amplicon sequence variants (ASVs) with feces from animals (7.4%), water (2.2%) and food (3.1%). Sharing of ASVs between humans and companion animals was not associated with household. There is a correlation between an Enterobacteriaceae ASV and the presence of blactx-m-2 in feces from humans and animals, hinting towards an exchange of antimicrobial resistant strains between reservoirs.

## 1. Introduction

Microbiota are complex and diverse ecological communities of microorganisms that are specific to a particular habitat such as specific human or animal body sites (e.g. the intestinal tract) or natural environments (1, 2). The native microbial composition is not only strongly associated with host factors but also with geographic and environmental variables (1, 3). Physical interactions between hosts and their natural environment may also alter the microbiota diversity and composition. Recent work on the microbiota of humans and animals suggested that due to interactions, such as contact with natural environments and co-housing conditions, individuals may share specific bacterial strains or have a more similar microbiota with each other and their environments (4, 5). Insights into the interconnections between the human and their environmental microbiome are important for understanding the dynamics in bacterial communities (5, 6).

Previous small-scale studies on specific populations demonstrated that antibiotic use is a key factor resulting in diversity loss of the human and animal microbiota. Furthermore, antibiotic pollution into natural environments influences soil and aquatic microbial ecosystems as well (6-8).

A study conducted among twelve healthy volunteers Caucasian males aged between 18 to 45 years showed that treating healthy individuals with meropenem, gentamicin and vancomycin led to an enrichment of antibiotic resistance genes (ARGs) and loss of butyrate producers species (9). In addition, repeated exposure to antibiotics not only drove the microbiome into a new steady-state that was different from the original state but also increased the survival and colonization potential of antibiotic resistant bacteria [11-13]. To date, most research on the effects of antibiotics has, however, been focused on microbial perturbations among study subjects without correlation to other co-habitats (10). To better understand host-microbiota relationships, it is important to put the research of the microbiota in the broader environmental-ecological context (11). Furthermore, it is of interest how microbiota adapt in settings where antibiotic exposure is expectedly frequent. Few studies have made such an attempt to characterize and compare the gut microbiota among individuals and their environment in a community setting with high antibiotic use (10, 12).

As an agricultural country, with nearly 50% of the population consisting of farmers in rural areas according to the report of general statistics office in 2019, Viet Nam has a history of using antibiotics in treatment and prevention in humans, animals and aquaculture (13–15). The impact of antibiotic use in Viet Nam on both community and hospital settings has been evaluated in many different aspects, including economic impacts, water pollution as well as the diversity and abundance of the resistome (16–18). However, there are still large gaps in our understanding of the human microbiota and its interactions with non-human microbial ecosystems. Here, we used 16S ribosomal RNA gene amplicon sequencing to characterize the microbiota of a cross-sectional collection of stool samples from humans and their domestic animals, and their food and water taken from a rural community cohort in Ha Nam, Vietnam. We describe the microbial diversity, composition and community structure of feces from humans and domestic animals and their environment in association with antibiotic consumption and high background of antibiotic resistance genes.

## 2. Results

### 2.1. Sample types and their microbiome sequence information

The V4 hypervariable region of 16S rRNA gene was sequenced for taxonomic analysis using QIIME2 to examine and compare the microbial compositions in human and animal feces, processed food and water. After exclusion of samples with low sequencing depth (n=4), 107 human fecal, 36 domestic animal fecal, 89 water and 74 food samples were retained for downstream analysis. We obtained 3,061 ASVs after clustering of 25,543,716 high – quality reads from 306 samples.

### 2.2. Microbiota of human gut in the study cohort

#### 2.2.1. Composition of gut microbiota of individuals in the context of demographical variables of study cohort

Overall, the human gut 16S rRNA gene sequences could be assigned to 13 phyla. The top four dominant phyla, accounting for 98.9% of all sequences, were *Firmicutes* (59.6%), *Bacteroidetes* (21.1%), *Actinobacteria* (9.3%) and *Proteobacteria* (8.8%). The most abundant genera (Figure 1) were *Faecalibacterium* with a median relative abundance (RA) of 5.4% [IQR 2.2 – 8.1], *Blautia* (5.2% [3.5 −7.9]), *Prevotella* (5.1% [0.5 - 20.3]), and *Bifidobacterium* (1.76% [0.0 - 6.6]) (Figure 1).

**Figure 1.**
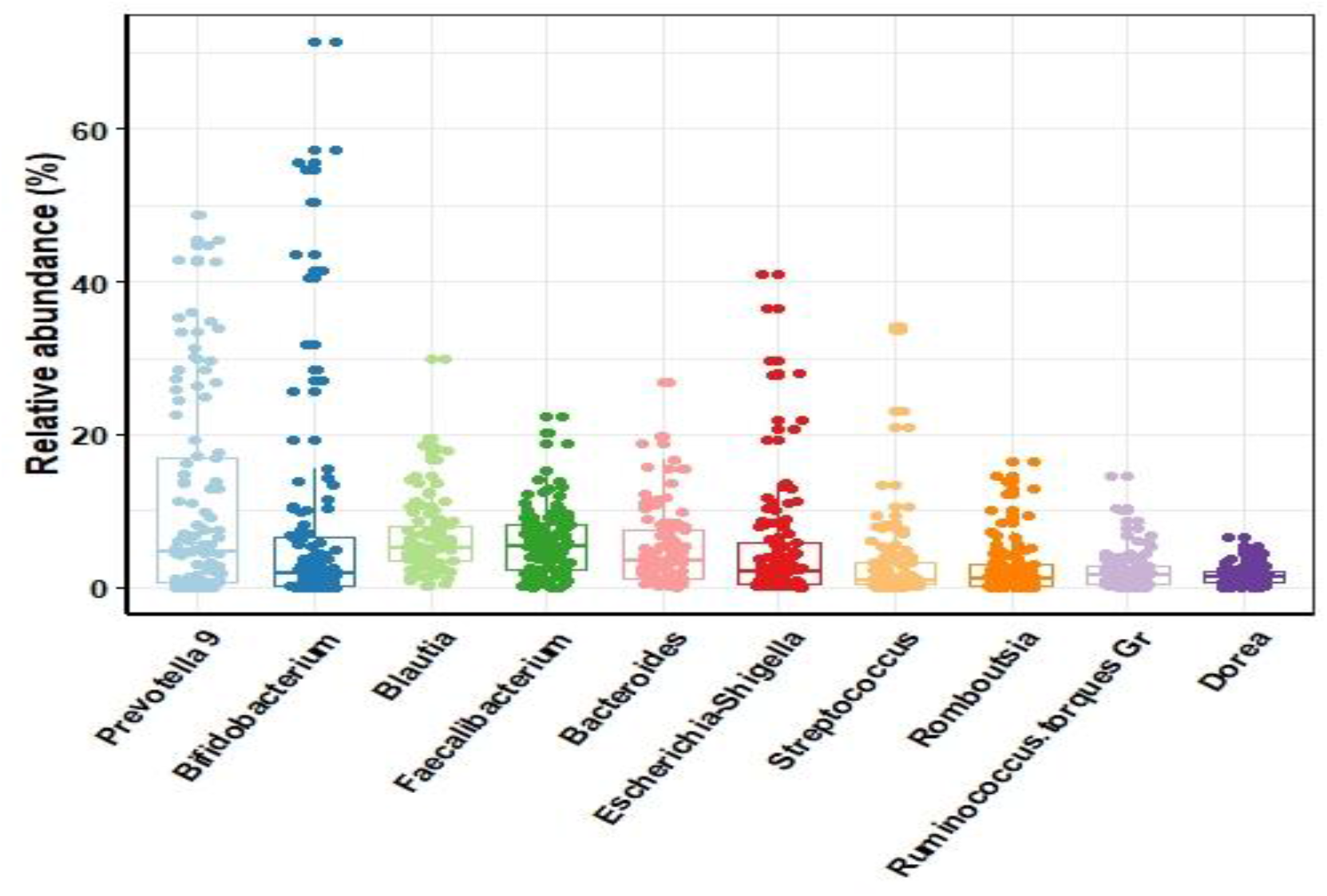
Relative abundance of the most predominant genera in human faecal samples (n=107). Boxes show the median and interquartile ranges, while whiskers show the 95% ranges.

To assess the compositional profile of the human gut microbiota, we compared the microbial richness, estimated by the Chao 1 index, and the bio-diversity, assessed by Shannon index, as well as the microbial community richness that incorporated phylogenetic relationship between ASVs (Faith’s PD) in relation to age, gender, and geographical location (see supplementary – Figure S2, S3, S4, respectively). While there were no differences in composition of gut between male and female individuals in our study cohort, the gut microbial diversity of children aged 6 years and below was less diverse than that of older children and adults. Both the estimated richness (Chao1) and phylogenetic diversity (Faith’s PD) were significantly lower in the youngest age-group as compared to individuals in the age between 7 to 55 years (two-sided Wilcoxon test, FDR q=0.02 and q=0.03, respectively). In line, the microbial composition of children aged 6 years or below was significantly different from other subjects (Bray-Curtis q=0.006, unweighted-Unifrac, q=0.015). The significant difference in microbial community structure between age groups was an indicator to investigate which genera were differentially abundant. Comparing the relative abundances at genus level of individuals at three age groups (0-6 years (n=17), 7-54 years (n=79) and ≥55 years and over (n=11) (Figure 2a) without additional adjustment for other risk factors, we observed a gradual decrease of bifidobacteria with age (ANCOM, W =161). In addition, the relative abundance of *Sellimonas* belonging to the Lachnospiraceae family, in children aged 0-6 years was relatively higher than other age groups (ANCOM, W=136) (Figure 2b). These observations withstood adjustment for antibiotic use and geographical region (ANCOM, W= 174 and W=167, respectively).

**Figure 2.**
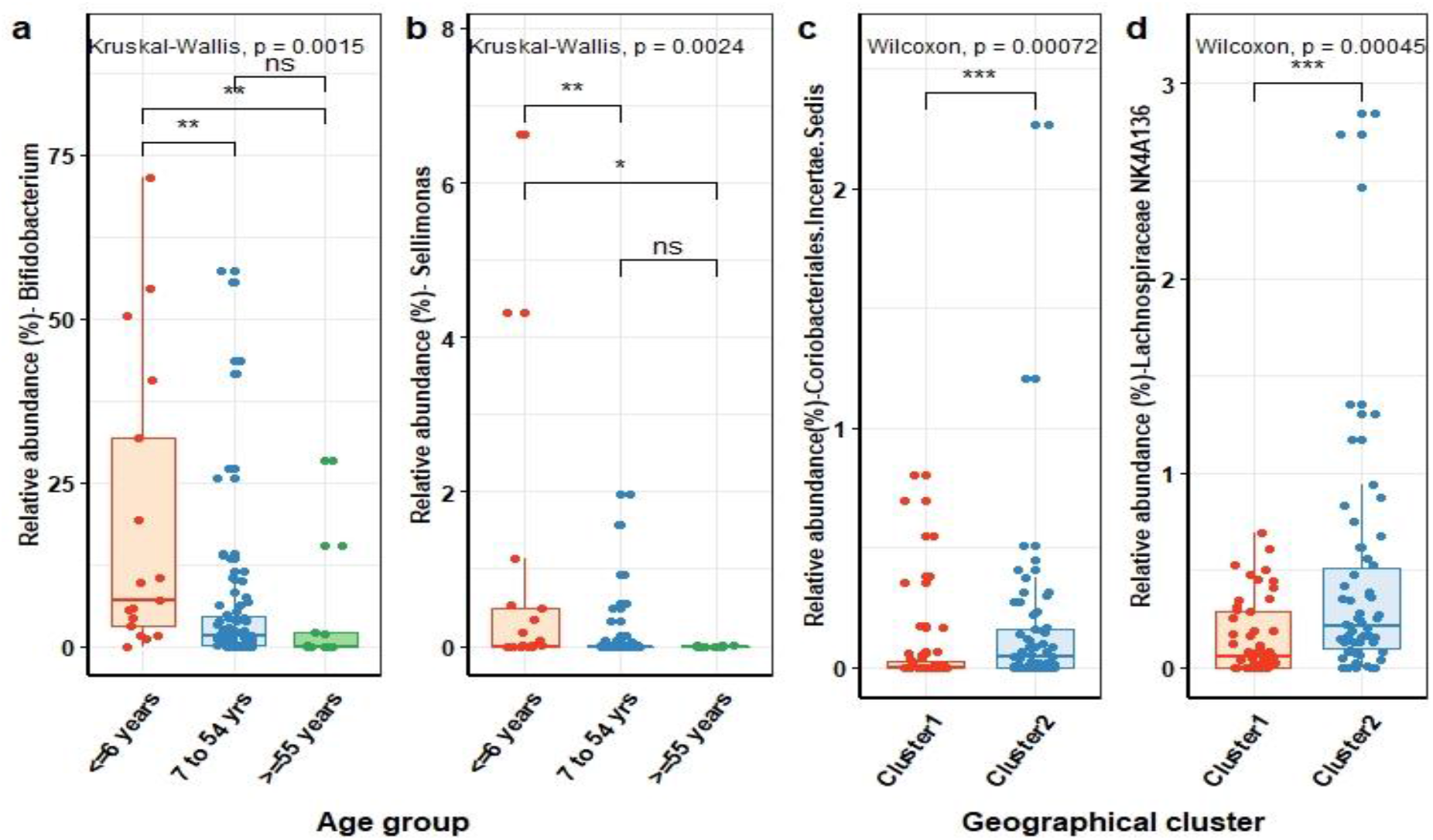
Human gut microbial genera in association to age (0-6 years (n=17), 7-54 years (n=79) and ≥55 years and over (n=11)) and geographical location cluster 1 (n=49) and clusters 2 (n=58). Comparison of relative abundance (%) of Bifidobacterium (a) and Sellimonas (b) genera between age groups. Comparison of relative abundance (%) of *Coriobacteriales Incertae Sedis* (c) and *Lachnospiraceae NK4A136* (d) between individuals living in the two geographical clusters.

We next examined the microbial composition of individuals in the context of living in geographically distinct sub-settings. Although the individuals lived in a narrow geographic area with similar living conditions and culture, we observed a significant lower microbial richness and biodiversity (two-sided Wilcoxon test, FDR q=0.014 and q=0.0009, respectively) among individuals living in cluster 1 (n=49) as compared to individuals in clusters 2 (n=58) (see supplementary - Figure S4). Comparing the microbial community structure of the human gut microbiota demonstrated quantitative differences between individuals living in cluster 1 and those living in cluster 2 (Bray-Curtis, q=0.03). Meanwhile, no differences in qualitative dissimilarities in microbial community structure were observed (unweighted-Unifrac, q=0.135), indicating that the majority of species are not specific to the gut microbiota of individuals living in a specific region but that some species may rather differ in their abundance. We indeed observed that the abundance of *Coriobacteriales Incertae Sedis* (ANCOM, W=119) family and *Lachnospiraceae NK4A136* (ANCOM, W=132) genus was significantly higher in the gut microbiota of individuals living in cluster 1 compared to individuals in cluster 2 (Figure 2c, 2d). These associations withstood adjusting for antibiotics use and age.

Examining the shared bacterial taxa between individuals in the same and in different geographical cluster showed that individuals living in cluster 2 share more ASVs with each other than with individuals from cluster 1 (27.2% [20.3 - 32.8] versus 26.0% [19.5 - 31.3], p = 6.83E-05). However, individuals in cluster 1 did not share more ASVs with each other than with individuals from the opposite cluster. This suggests that there is more dispersal of bacteria among individuals within cluster 2.

As dispersal of bacteria is likely most prominent within households, we next examined to what extend individuals within the same household share more bacterial taxa than individuals from different households. These results confirmed that the proportion of shared bacterial taxa was significantly higher among individuals from the same household (median 27.9% [21.3 - 34.1] as compared to individuals from different households (26.2% [19.5 - 31.6], Wilcoxon p = 8.05*10^− 3^). When stratifying these analyses for age, both children under the age of 6 years, as well as individuals above the age of 6 years, share more bacterial taxa when living in the same household (see supplementary Table S3). These results thus confirm the sharing of bacteria among household members.

#### 2.2.2. Composition of gut microbiome of healthy people in the context of their antibiotic use history

Among all 107 participants, 43 individuals used antibiotics at least once during the past 4 months. Like previous findings about the influence of antibiotic use on microbial compositions, we observed a reduction in gut microbial richness and diversity among individuals who used antibiotics (n=43) as compared to individuals who did not (n=64) use antibiotics in the past 4 months (Figure 3a). Both the estimated richness (Chao1) (Figure 3a) and phylogenetic diversity (Faith’s PD) (Figure 3c) were significantly lower in individuals who used antibiotics (two-sided Wilcoxon test, P value q=0.007 and q=0.017, respectively), whereas the microbial diversity as measured by the Shannon index did not reach statistical significance (two-sided Wilcoxon test, P value q=0.08) (Figure 3b). In addition, Aitchison distance showed significant differences in microbial community structure in association to antibiotic use (PERMANOVA, q=0.009). The results were confirmed when using other beta-diversity indices that measure not only the dissimilarities between microbial composition but also incorporate phylogenetic relationships (unweighted Unifrac, q = 0.016). However, no clear visual separation in the microbial community structure of individuals with or without recent antibiotic use could be observed when visualizing these dissimilarities using principal component analysis (PCA). This indicates that although recent antibiotic use significantly influenced the microbial community structure, it did not result in profound shifts (Figure 3d). We next investigated which specific taxa were perturbed due to antibiotic use in our cohort. Unlike previous studies, we observed only few genera, belonging to *Clostridiales* order, to be significantly associated with antibiotic use (two-sided Wilcoxon test q<0.05, antibiotic use variable without adjusting for age and geographic variables), including *Lachnospiraceae FCS020 group*, and *Ruminococcus gnavus group* genera. The relative abundance of *Lachnospiraceae FCS020 group* in individuals who used antibiotics in the past 4 months was significantly lower than in individuals without antibiotics use (ANCOM, W=124) and this difference withstood adjustment for age and geographic variables. When adjusting for either age or geographic variables the difference in relative abundance of the *Ruminococcus gnavus group* was no longer statistically significant but we did observe a slight decrease in relative abundance of *Ruminococcaceae UCG-002* in individuals who used antibiotics in the past 4 months as compared to individuals without antibiotic use (ANCOM, W=120) (see supplementary Table S4).

**Figure 3.**
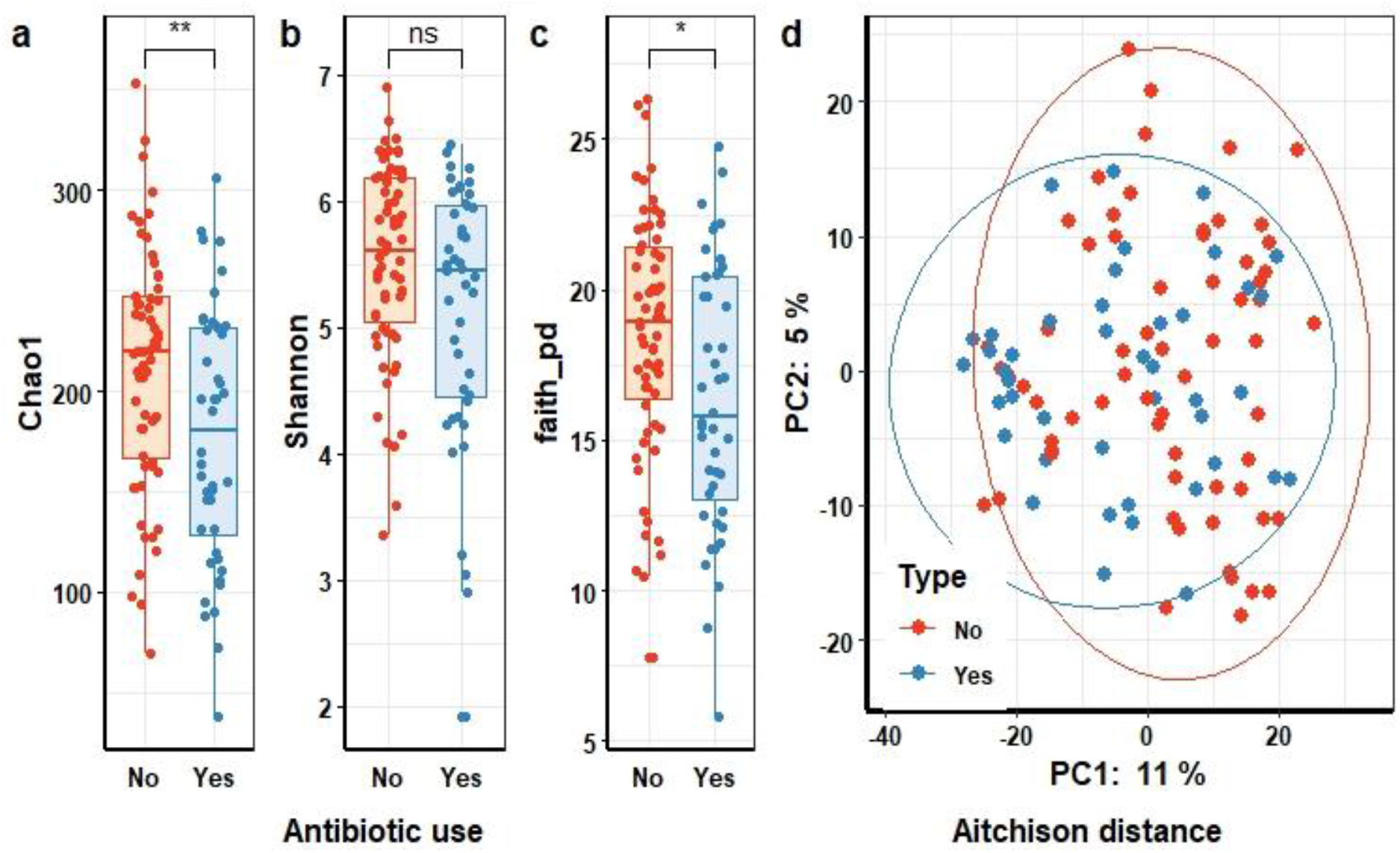
Differences in gut microbial community structures and taxonomic composition in association to antibiotic use in the 4 months prior to faecal sampling. Differences in microbial richness (Chao1) **(a)** and diversity (Shannon **(b)** and Faith’s PD **(c)**) between people who did (n=43) or did not use (n=64) antibiotics. Statistical significance (P-value<0.05) determined by Wilcoxon test. **(d)** Principal Component Analysis based on Aitchison distance colored according to antibiotic use (PERMANOVA test, P-value<0.05). Each point represents the gut microbial community structure of an individual that did (blue) or did not (red) use antibiotics.

### 2.3. Microbiota of non-humans samples

#### 2.3.1. The microbial composition of domestic animals

To perform in-depth characterisation of the non-human microbiomes in the present community, we investigated the differences in composition between sample types (i.e. animal feces, food and water) as well as between samples from the same origin but from the different geographic regions. Interestingly, we neither observed differences in microbial richness and biodiversity nor in microbial community structure between the two domestic animal species, chicken (n=11) and dogs (n=24) (see supplementary Table S5). When comparing alpha diversity values between animals living in different geographical clusters, the microbial richness and biodiversity of animals in cluster 1 (n=21) was significantly higher than in cluster 2 (n=14) (two-sided Wilcoxon test; q<0.05) (Figure 4a,4b, 4c). Although we observed no dissimilarities between cluster 1 and cluster 2 in measures of microbial community structure (PREMANOVA, q >0.05) (Figure 4d), the qualitative dissimilarity, incorporating phylogenetic relationships, appeared significantly different (unweighted UniFrac distance, PERMANOVA q=0.029 - see supplementary Table S5). These findings indicate that the microbial community structure of feces from domestic animals was associated with the geographical location and could be related to the contamination from soil, as we could not collect the fresh feces from domestic animals.

**Figure 4.**
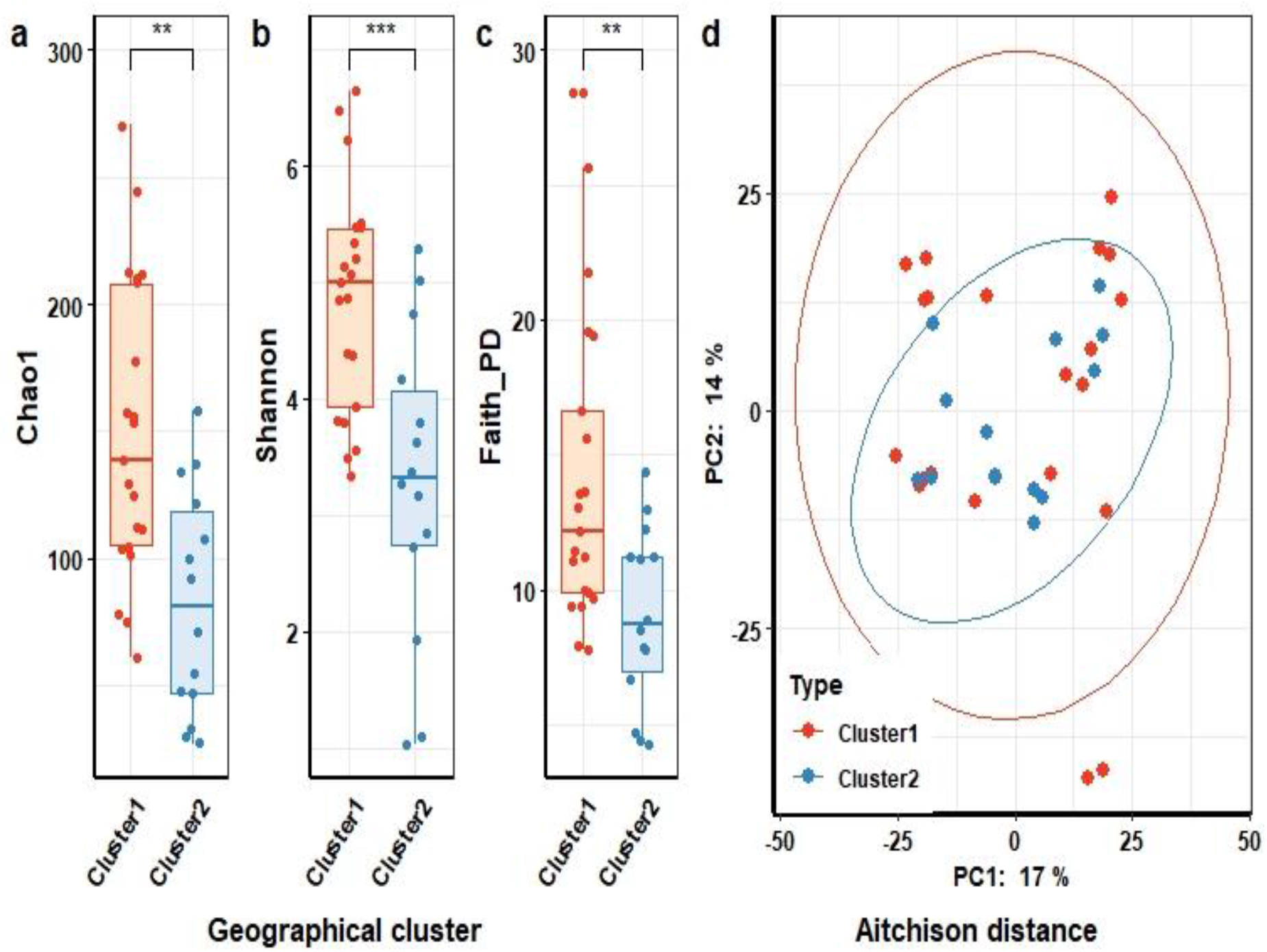
Microbial richness, diversity and community structure of the chicken and dog gut microbiota. Differences in the microbial richness (Chao1 **(a)**) and diversity (Shannon **(b)** and Faith’s PD **(c)**) between chickens (n=11) and dogs (n=24) compared by Wilcoxon test, difference is significant at P-value<0.05. **(d)** Ordination of the gut microbiota community structures of geographical cluster based on Principal Component analysis of Aitchison distance. Each point represents the gut microbial community structure of an individual animal and is colored according to geographic cluster with animals from cluster 1 (n=21) colored in red and animals from cluster 2 (n=14) colored in blue. (PERMANOVA test, P-value>0.05 for geographic cluster).

We also investigated which microorganisms were dominant and more abundant in different animal species. At genus level, 181 genera, accounting for 98.3% of total sequences, were detected across all samples from domestic animals. The most dominant genera were *Escherichia* with a median of relative abundance (RA) of 7.1% [2.0 −15.6]), *Lactobacillus* (median 5.5% [0.5 – 18.8]), *Acinetobacter* (median 3.2% [0 – 10]) and *Clostridium sensu stricto* (median 2.3% [0.5- 6.0]). Significant differences in relative abundance at genus level were however neither observed when comparing different animal species nor when comparing animals from the different geographic clusters. These latter results should however be interpreted with caution due to the potential lack of statistical power related to the relatively small sample size.

#### 2.3.2. The microbial composition of water

Unlike the findings in domestic animals, water in different sources have different compositions. The estimated microbial richness (Chao1), Shannon and phylogenetic diversity (Faith’s PD) all significantly differed between types of water, with irrigation water being the most diverse, followed by rainwater and water from wells (two-sided Wilcoxon singed-rank test; q<0.05) (Figure 5a, 5b, 5c). PCA demonstrated that the microbial composition of irrigation water was profoundly different from the composition of the other two water sources (Aitchison, PERMANOVA, q=0.001) (Figure 5d), and this held true for other indices of microbial community structure (Bray-Cutis, unweighted UniFrac, PERMANOVA, both q=0.001) (see supplementary – Table S6). We identified 28 phyla in water samples from three sources, which is far greater than the 13 and 17 phyla observed in humans and animals, respectively. At a lower taxonomic level, among the most prevalent 349 genera, which were detected in at least 10% of all water samples (n=89), accounting for 98.1% of total sequences, there were 184 uncultivable or unidentified genera, accounting for 41% of total sequences. The genus *Acinetobacter* and *Burkholderiaceae* family were the most abundant bacteria with median of RA at 5.4% [0.5 – 38.4] and 6.6% [3.2 - 16.4], respectively. *Novosphingobium, Sediminibacterium, Pseudomonas* and *Aquabacterium* genera were detected at low abundance (ranging in median from 0.2% [0.1 – 1.1] to 0.9% [0.2 – 4.2]), but each of these genera was highly prevalent and detected in more than 90% of all water samples (Figure 5e). The difference in microbial community structure between water sources was also evident when examining the relative abundance of specific microbial taxa. Overall, 116 genera differed in abundance across the water samples from different sources (ANCOM, W>231), see supplementary-Table S7). These differences strongly supported that the water’s source was the key factor responsible for the distinction and unique compositional profiles between water samples.

**Figure 5.**
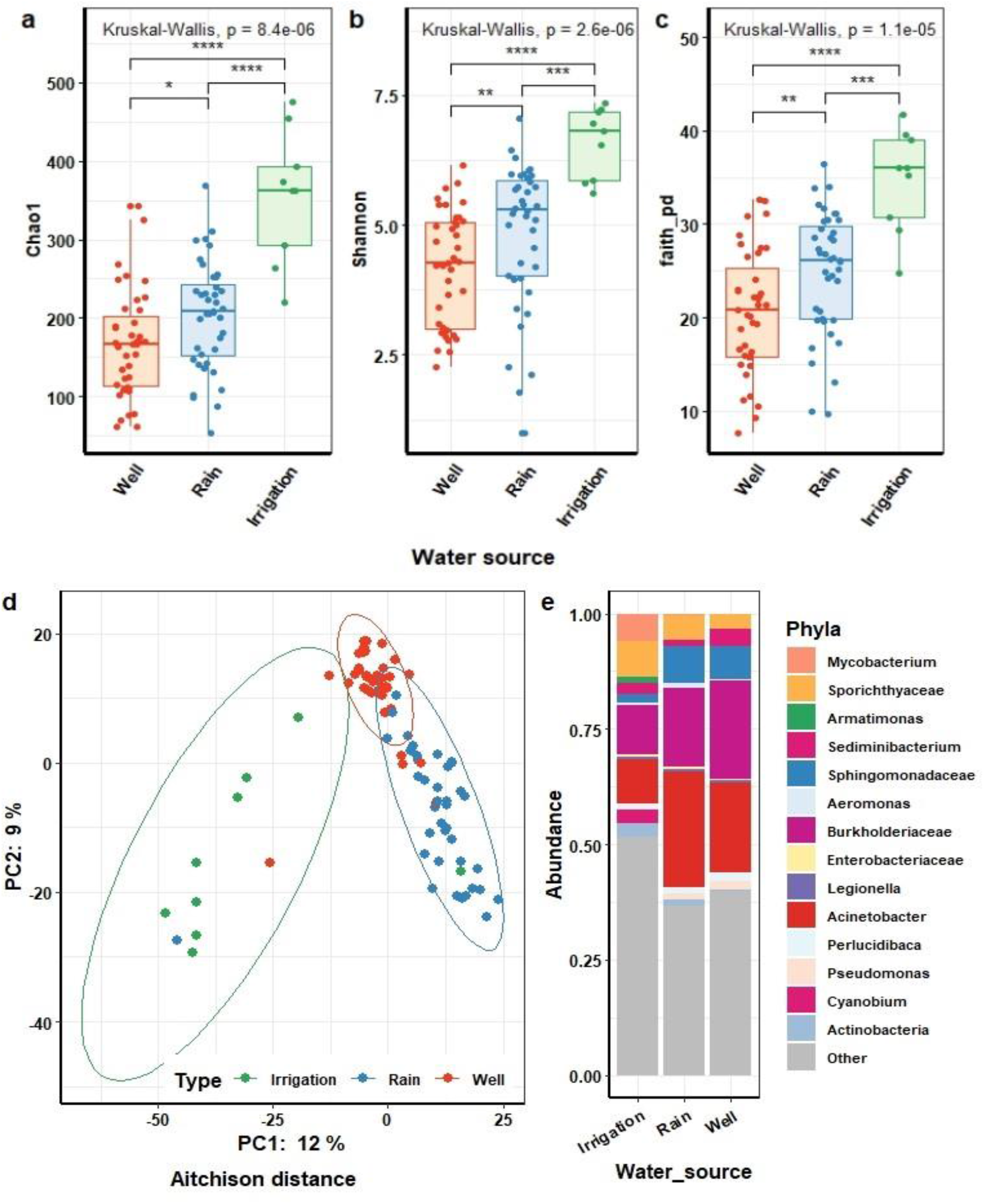
Microbial richness, diversity, community structure and dominant taxa of different water sources (n=89). Differences in microbial richness (Chao1) **(a)** and diversity (Shannon **(b)** and Faith’s PD **(c)**) of different water sources compared by Wilcoxon tests. Difference is significant at P value<0.05. **(d)** Ordination of the microbiota community structures of water **(Figure 5 – continued)** samples based on Principal Component analysis of Aitchison distance (PERMANOVA test, P value<0.05). Each point represents the microbiota community structure of well (red) (n=40), irrigation water (green) (n=9) and rainwater (blue) (n=40), respectively. **(e)** The relative abundance of major bacterial genera in different water sources.

#### 2.3.3. The microbial composition of processed food

In this study, we also investigated the microbiota of cooked food including food that individuals eat in their daily meal at a single time point. We observed that the top four phyla, identified in at least 83.8% of food samples, were Firmicutes (75/75), Proteobacteria (74/75), Bacteroidetes (63/75) and Actinobacteria (62/75). In our study design, processed food at single time point has been considered as a low biomass microbiota and the composition is likely not representative for the overall microbiota related to the food consumed by the participants in our study cohort. Therefore, we only used these data as an additional non-human microbiota in order to investigate the potential dispersal of taxa between individuals and their environmental factors.

### 2.4. Human versus non-human microbiota

#### 2.4.1. Overall composition and diversity of the human gut microbiota in relation to microbial community of domestic animal gut, water and processed food

We explored the microbial composition of the entire cohort and visualized the shared microbial taxa between humans and non-human samples. The microbial richness and diversity of human fecal samples was significantly higher as compared to feces from animals and food samples, but not of water from different sources, in our cohort. Both estimated richness (Chao1) and Shannon diversity were extremely high in human as compared to animal and food samples (two-sided Wilcoxon, FDR-adjusted P-values (q) = 1.472e-07 and q=4.421e-06, respectively for humans versus animals and FDR-adjusted P-value (q) = 5.343e-25 and q=3.051e-24, respectively for humans versus food). In addition, results were similar when assessing community richness by including phylogenetic relationships between ASVs using Faith’s PD. Despite no significant difference in estimated richness (FDR-adjusted P-value q = 0.75) between human and water (Figure 6a), the Shannon diversity, which incorporates both abundance and evenness of taxa, and the phylogenetic diversity (Figures 6b, 6c), remained significantly higher in human as compared to water samples (two-sided Wilcoxon, FDR-adjusted q=5.56e-04, q=1.14e-09, respectively). In line with significant differences in alpha diversity, PCA demonstrated that human gut microbiota clustered apart from domestic animals, water and food along the first component. Meanwhile domestic animals, water and food were relatively more similar to each other (Figure 6d).

**Figure 6.**
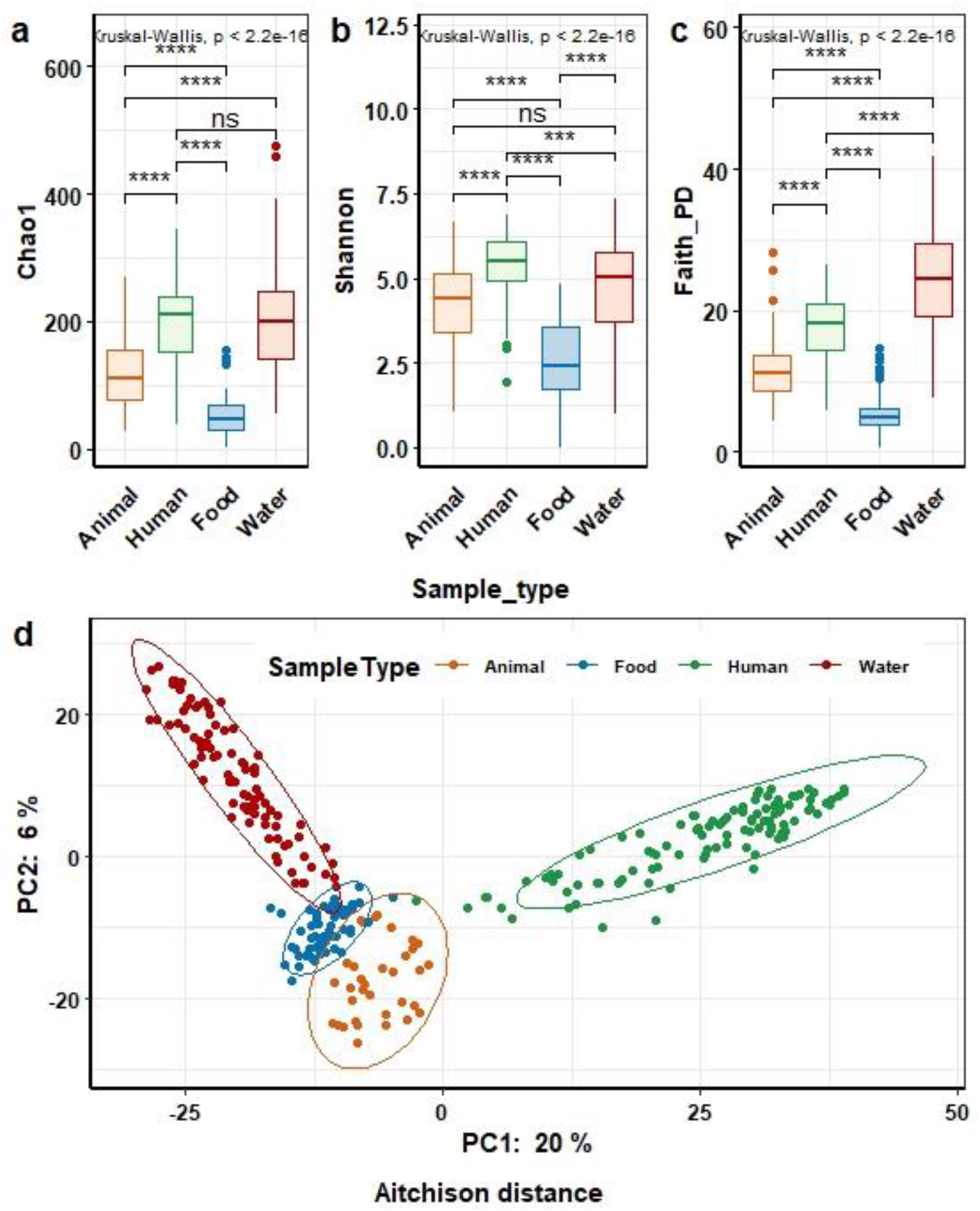
Microbial composition of human and animal gut, water and food.Differences. **in e**stimated microbial richness (Chao1) **(a)** and diversity (Shannon **(b)** and Faith’s PD **(c)**) across sample types. Kruskall-Wallis test followed by Wilcoxon test for post-hoc comparisons between four sample types (Humans (n=107), Animals (n=35), Food (n=75) and Water (n=89)). ***P<0.001, ****P<0.0001. **(d)** Principal Component Analysis (PCA) based on Aitchison distance. Each point represents the microbial community structure of an individual animal stool (orange), human stool (green), food (dark blue) and water (red) sample. (PERMANOVA test, P value <0.001).

The microbial community composition of human and animal gut, processed food and water was assigned to 31 bacterial phyla. The top seven dominant phyla, accounting for 98.7% of the total sequences, were *Firmicutes* (38.8%), *Proteobacteria* (38.7%), *Bacteroidetes* (11.9%), *Actinobacteria* (6.7%), *Planctomycetes* (1.1%), *Verrucomicrobia* (0.9%) and *Fusobacteria* (0.3%). The *Firmicutes* phylum was predominant in the human (median relative abundance 62.0% [46.5-73.3%]) and animal (median 53.2% [39.4 - 74.3%]) gut and food (median 47.8% [11.8 –91.6%]). Meanwhile, *Proteobacteria* was the primary phylum in water (median 72% [58.0 – 87.0]), while also being highly abundant in food (median 47.7% [7.2 – 85.0]) and in animals stools (median 29.4% [6.3 – 44.3]). The *Verrucomicrobia* phylum was low abundant in all sample types with a median relative abundance ranging from 0 % [0 – 0.001] in feces from human to 0.9 % [0.1 – 3.5] in water, however this phylum was much more prevalent in water samples (96%, 87/91) than in human feces (32.7%, 35/107).

At lower taxonomic levels, 524 genera, accounting for 99.3% of total sequences, were detected across 306 samples. The genera *Acinetobacter, Prevotella, Staphylococcus*, and genera belonging to family *Enterobacteriacea* were the most abundant, accounting for 27.2% of all sequences. Many genera were presented in all four types of samples such as *Streptococcus, Bacillus, Comamonas, Lactococcus* and *Acinetobacter* (Figure 7). Meanwhile some other genera were only prevalent in specific sample types such as *Mycobacterium* and *Fluviicola* in water, *Intestinibacter* and *Roseburia* in human gut. Notably, our study showed that many genera belonging to *Proteobacteria* and *Actinobacteria*, were detected in both animal gut and water but not or rarely in the human gut.

**Figure 7.**
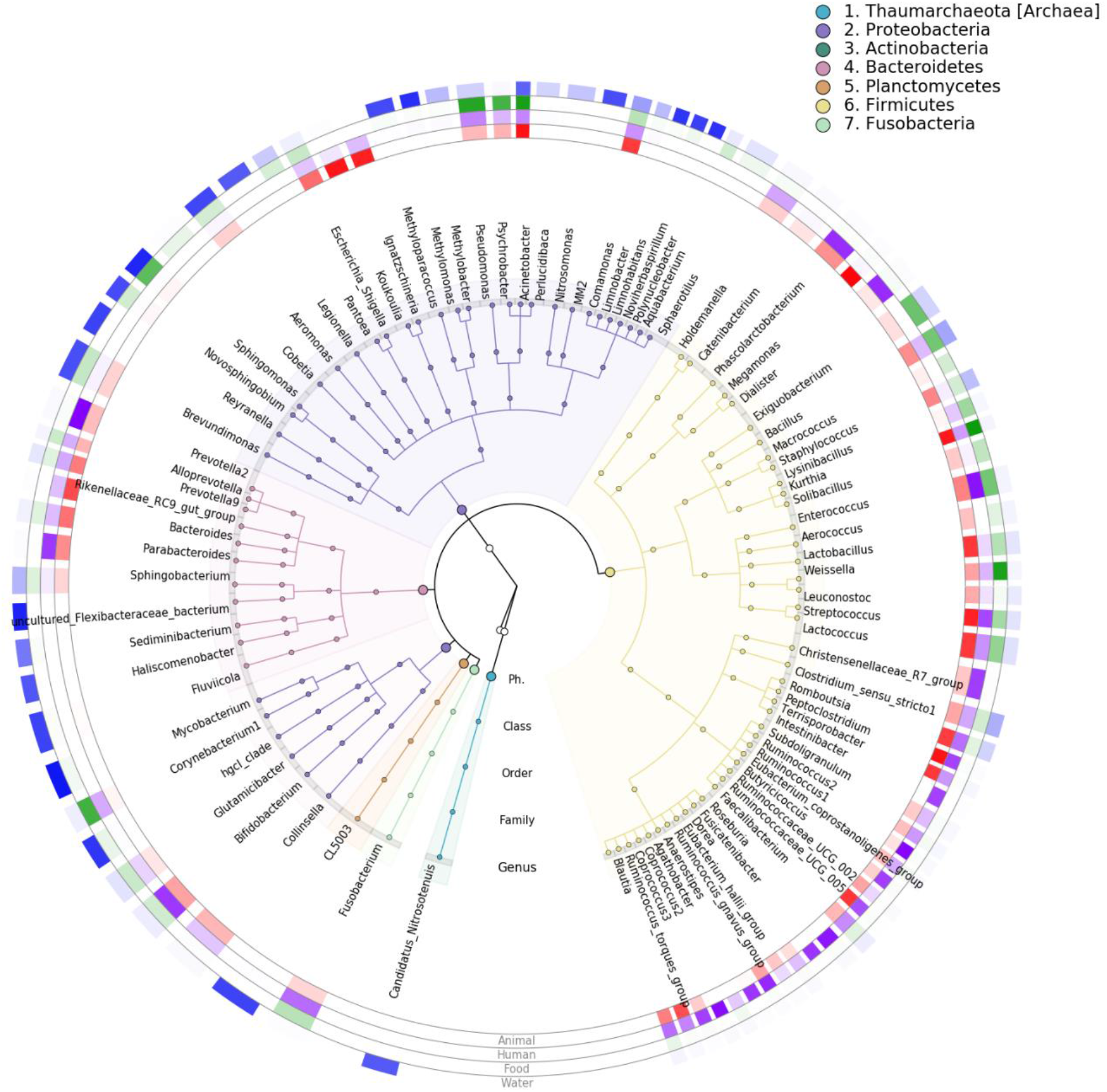
Cladogram depicting the relative abundance of the bacterial genera detected in the different sample types. Only genera with a mean relative abundance of >0.5% in at least one of the sample types were included. Background and branch colors reflect the different phyla. The colour density of the four outer rings reflects the relative abundance (arcine square root transformed) of the genera in the different sample types (with opaque color indicating a relative abundance of 1 and fully transparent indicating a relative abundance of 0).

#### 2.4.2. Potential dispersal of microbial taxa between the microbiota of humans and non-humans

Although the depth of strain-level cannot be achieved with 16S rRNA gene amplicon data, amplicon sequence variants give a much better resolution than clustering into operational taxonomic units. We therefore explored the potential dispersal of ASVs between microbial ecosystems. We observed that humans shared on average 26.2% [19.6 – 31.6] ASVs in their feces with other individuals. When comparing human feces with other microbial habitats, we observed that feces from humans shared a median of 7.44% [3.2 – 9.9] of its ASVs with the feces from domestic animals, 2.2% [1.2 – 2.8] with water and 3.1% [1.5 – 4.1] with processed food. The proportion of shared ASVs between human and non-human samples from the same HH was however not significantly higher than the proportion of shared ASV’s between those samples from different HHs (see supplementary Figure S5). The most commonly shared genera between humans and non-humans belonged to the Enterobacteriaceae family.

In order to investigate specific ASVs within genera that are widely shared between humans and their living environment, we subsequently focused on those ASVs classified as Enterobacteriaceae. We observed that 50 ASVs classified as Enterobacteriaceae across sample types. Each individual shared at least one Enterobacteriaceae ASV with animals, water and food samples, however, the proportion of these shared ASVs between human and non-humans samples did not significantly differ between samples from the same as compared to samples from different HHs (Wilcoxon, p-value >0.05) (see supplementary Figure S6). Interestingly, when looking at the ASVs shared between different hosts and associated with specific ARGs, we observed several ASVs that were positively correlated with the presence of ARGs in both human and animal samples suggesting exchange of antimicrobial resistant strains between human and animal reservoirs. In particular, we observed that an ASV belonging to the Enterobacteriaceae family and present in 46.7% (50/107) of human samples and 28.6%% (10/35) of animal samples was positively correlated with the presence of *bla*_*ctx-m-2*_in faecal samples. In 30.0% (15/50) of humans carrying this ASV and 40% (4/10) of animals carrying this ASV we also detected *bla*_*ctx-m-2*_ (p = 0.0016 and p = 0.026, respectively).

## 3. Discussion

In this study, we characterized the microbiota of the human and domestic animal gut and their associated environments using 16S rRNA gene sequencing in a community setting with a high frequency of antibiotic consumption. We described the core group of bacterial taxa in feces from humans and domestic animals, food and water from three different sources. The results of our study indicated that antibiotic use decreased the microbial diversity in the human gut, although only few bacterial taxa appeared to be affected in the present population. We furthermore showed that even in the confined habitat of seven neighboring villages, the microbiota composition of both humans and animals appeared to be associated to geographic location. Finally, the human gut microbial composition differed most from all other sample types analyzed in the present study and the limited amount of shared ASVs between habitats indicates the presence of barriers for microbial dispersal between ecosystems.

In the current study, we focused on describing the patterns of human gut microbiota in the cohort with high level of antibiotic use. Our results showed that the *Prevotella, Bifidobacterium, Blautia, Faecalibacterium* and *Bacteroides* genera were the most abundant among rural individuals in Vietnam. The median relative abundance of *Bifidobacterium* in our population was 1.76% [0 – 6.6] which is similar to the estimated abundance of *Bifidobacterium* of below 2% as described in US, South Korea, and Europe (19, 20). The *Bifidobacterium* genus is known to confer health benefits through metabolic activities such as involving in oligosaccharide fermentation, but the increase of its relative abundance in human adult gut composition is an indicator of antibiotic use such as amoxicillin, ampicillin and gentamicin as described previously (21, 22). Interestingly, we observed a higher abundance of the Proteobacteria phylum (median 3.5% [1.3 – 9.0]) when compared to Europeans, Americans and Asians in other areas (less than 2%) (23). Especially, it was much higher than that of individuals living in other traditional populations with low antibiotic exposure such as the Hadza hunter-gatherers of Tazania or the Yanomami in the Amazon jungle (24, 25). We found that the Enterobacteriaceae family consists mostly of pathogens with a high level of antibiotic resistance genes (26). Moreover, a previous study showed that there was an acute blooming of *Klebsiella spp* and *E*.*coli* in gut microbiota after antibiotic treatment (7). The relatively high abundance of Proteobacteria in the human gut microbiota in our cohort is likely the result of intensive exposure to antibiotics (27).

Our results are in agreement with recent studies indicating the significant relationship between age groups and the abundance of particular genera (28–30). For instance, our finding was consistent with previous reports of the decreasing abundance of bifidobacteria with age (31). The *Sellimonas* genus was also found to decrease with age but there is limited evidence on its critical role in the gut microbiota so far (32). In this study, we indicated the difference in the relative abundance of the *Lachnospiraceae NK4A136* genus between individuals living in two geographical locations. In previous reports, the low abundance of this genus, which belongs to short-chain fatty producing bacteria, was associated with obesity in humans and mice (33–35). We only recorded that the households living in cluster 1 consumed meat more frequently than the households living in cluster 2 (more than once versus one time per week, Chi-square, p value <0.05), but we had no information to link the abundance of this genus to the body mass index of individuals.

In contrast to the majority of previous studies (7, 36), we observed only a modest difference in microbial composition between individuals who used and who did not use antibiotics in the previous four months. Previous studies reported that after short-term antibiotic use, the diversity of the gut microbiome is dramatically reduced and needs several months to recover to its original state. Some other studies demonstrated that many common species in the gut microbiome of healthy adults disappeared up to 180 days after antibiotic treatment (7, 37). Additionally, we only observed that the abundance of *Lachnospiraceae FCS020 group* was significantly reduced among individuals who recently consumed antibiotics. Our results might suggest that frequent exposure to antibiotics in the entire population might have resulted in a new steady state of microbial composition that is less influenced by subsequent antibiotic pressure (36, 38). This slight impact might also be explained by the fact that beta-lactam antibiotics, whose perturbations on the microbiome are less profound than macrolides or quinolones, were the most commonly used in our study.

Our results also highlight the similarities and dissimilarities in the microbial community structure between human and non-human samples. We observed that human gut had the highest diversity followed by water, domestic animals and cooked food by considering the estimate richness and biodiversity indices. *Firmicutes* and *Bacteroidetes* consist of major genera, which play critical roles in forming the gut microbial composition of humans and animals (2, 28). In contrast, the phylum *Proteobacteria*, which included the genera *Pseudomonas, Acinetobacter* and *Enterobacteriacea* family, is the most predominant in water including freshwater, sewage and river (39, 40). Although the microbial compositions of humans, animals and water are unique, our results showed a small overlap at ASV level. In the same habitat, human gut microbiota shared nearly 8% of their ASVs with domestic animal gut and 4% with water. Most of shared ASVs between humans and non-humans samples were classified as *Enterobacteriacea* family, which is a potential reservoir for shedding antibiotic resistance genes. Recent studies suggest that through feces, humans and domestic animals disseminate antibiotic resistant bacteria into the environment (41). Our previous report showed that antibiotic resistance genes were detected in water at low proportion as compared to humans and animals (42). Along with the antibiotic pollution in water resources, antibiotic resistant microbes might become a part of water microbiota (43). Therefore, a possible consequence of the high pressure of exposure to antibiotics is that humans may become more receptive to be colonized with resistant bacteria from their environment.

The present study has several limitations. Due to the cross-sectional nature of the sample collection, we were not able to longitudinally follow the microbiota recovery after antibiotic exposure. Additionally, the 16S rRNA gene amplification-based sequencing approach did not allow us to directly match the antibiotic resistance genes (ARGs) with their carrier taxa. Using sequence-based metagenomics, a recent study linked ARGs with the 50 most abundant genera in urban sewage from 60 countries (39), while another report also showed that most of the ARG-carrying sequences in migratory birds originate from Proteobacteria (40). Future studies collecting repeated samples at different time points are needed to further extend our knowledge on the long-term impact of antibiotic use on microbial ecosystems in low-and middle resource settings.

## Conclusion

In conclusion, we characterized the microbiota of feces from humans, domestic animals and their direct environment in a Vietnamese community with high antibiotic use. Our results are in agreement with previous studies about the transition points in gut microbial composition with age as well as indicate some microbial patterns related to geographical location. In addition, the small difference between antibiotic users suggested that exposure to antibiotics might not play an important role in microbial perturbation in the cohort with high background of antibiotic use. We provide a proof of taxa that could be potential ARGs carriers and might spread to human’s direct environment through their interaction with human gut microbiota. Indeed, we found a potential exchange of *bla*_*ctx-m-2*_ gene, which is one of the most prevalent ESBLs between feces from humans and animals in this study.

## 4. Material and methods

### 4.1. Community setting, study design and samples collection

#### -Community setting

The demographics of this complete cohort have been described in our previous study, which was designed to explore the circulation of ARGs in humans, animals and their environment (42) (see supplementary-Table S1).

For the present study, a subset of 44 / 80 participating households was selected: 23 households in which at least one member used any antibiotic in the four months prior to sample collection and 21 households in which no antibiotic use was reported prior to sampling. We used feces from humans (n=107) and domestic animals (n=36), processed foods (n=74) and water samples (water used for cooking and/or washing and irrigation) (n=89) collected at a single time point (at the fourth month of study – M4) collected from these selected households (HHs) to analyze the microbiota using 16S rRNA gene amplicon sequencing.

In this study, samples were collected from 44 households (HHs) residing in seven villages within 8.1 square kilometers. Lifestyles in these villages are very similar with diets dominated by rice, vegetables and meat. The frequency of meat consumption was depended on their economic condition of HHs. To evaluate the geographical impact on the microbiota, particularly, in order to identify whether individuals living closer together shared more ASVs or not, we separated the 44 HHs into clusters based on their locations. We used partitioning clustering with k-medoids algorithm, which enable us to find out two clusters in 44 HHs based on the distance from HH’s locations (detected by their Latitude and Longitude) to the centre of each cluster. The number of participating individuals living in cluster 1 with 23 HHs and cluster 2 with 21 HHs were 49 and 58, respectively (see supplementary Figure S1 and supplementary Table S2).

### 4.2. Methods

#### DNA extraction and 16S –rRNA amplicon sequencing

Metagenomic DNA has been isolated as described previously (42). In brief, Repeated Bead-Beating (RBB) combined with column-based purification was used to extract DNA from human and animal fecal samples according to protocol Q of the International Human Microbiome Standards consortium [20]. For isolation of microbial DNA from water samples, 100 ml of collected water was filtered through 0.22 μm mixed cellulose esters membrane filters (Sartorius, Göttingen, Germany) to capture bacteria. One quarter of the filters were used for metagenomic DNA extraction using the QIAGEN DNeasy Power Water kit according to the manufacturer’s protocol. Upon isolation, DNA concentrations were determined using the Quan-iT PicoGreen dsDNA assay (Invitrogen, Carlsbad (CA), USA).

Amplicon libraries and sequencing was performed according to previously published protocols (44). Briefly, the V4 region of the 16S rRNA gene was PCR amplified from each DNA sample in triplicate using the 515 f/806r primer pair (45). Pooled amplicons from the triplicate reactions were purified using AMPure XP purification (Agencourt, Massachusetts, USA) according to the manufacturer’s instructions and eluted in 25 µl 1 × low TE (10 mM Tris-HCl, 0.1 mM EDTA, pH 8.0) and subsequently quantified by Quant-iT PicoGreen dsDNA reagent kit (Invitrogen, New York, USA) using a Victor3 Multilabel Counter (Perkin Elmer, Waltham, USA). Amplicons were mixed in equimolar concentrations to ensure equal representation of each sample and, were sequenced on an Illumina MiSeq instrument using the V3 reagent kit (2x 250 cycles).

#### Bioinformatics analysis

Raw reads from the 16S-rRNA amplicon sequencing were demultiplexed and quality controlled using analysis package QIIME2-2019.7 (Quantitative Insights Into Microbial Ecology 2-2019.7) (46). We used DADA2 pipeline for sequence quality control and feature table construction. We truncated sequences at position 200 of forward reads and at position 140 of reverse reads to remove low quality regions of reads while maintaining sufficient overlap between forward and reverse reads. The high-quality reads resulting from denoising and chimera-filtering steps were clustered into a table of Amplicon Sequence Variant (ASV). An ASV refers to single DNA sequences recovered from a high-throughput marker gene analysis upon removal of erroneous sequences generated during PCR and sequencing and is thereby a higher-resolution analogue of the traditional Operational Taxonomic Units (47). Samples with a sequencing depth below 5,000 reads were excluded from downstream analyses and all ASVs with a relative abundance less than <0.0001%were discarded. We further removed contaminant ASVs annotated as Mitochondria and Chloroplast and, generated a tree for phylogenetic diversity analyses by FastTree. Taxonomic assignment of representative ASVs was carried out with the RDP (Ribosomal Database Project) classifier based on Silva database (release: Silva 132).

We used QIIME2 diversity analyses to compute alpha and beta diversity metrics. Alpha diversity is the ecological diversity within a sample. The following alpha-diversity measures were calculated: Chao1 index (estimated microbial richness), Faith’s Phylogenetic Diversity (Faith’s PD, richness index that additionally incorporates the phylogenetic relationships between microbial taxa) and Shannon index (true biodiversity index not only taking into account microbial richness but also their evenness).

Beta-diversity is a measure to compare differences in microbial community structure between samples. Aitchison and Bray-Curtis distances were calculated as quantitative beta-diversity metrics (taking into account relative abundance profiles)(48), whereas the unweighted-UniFrac distance was calculated as a qualitative measure of community dissimilarity (presence/absence of microbial taxa) that incorporates the phylogenetic relationships between ASVs (49).

To investigate potential similarities in the microbiota of humans and non-humans, we determined the proportion of shared ASVs between all human subjects as well as between non-human samples and humans in the cohort. In addition, we identified the shared ASVs that were classified as Enterobacteriaceae family in order to trace the similarities between humans and their living environments.

#### Statistical tests

Non-parametric two-sided Wilcoxon-rank sum and Kruskal-Wallis tests were used to test whether species richness (Chao1, Faith’s PD) and diversity (Shannon) were significantly different between sample types and within individuals in relation to geographical location. These analyses were also performed to determine the effect of antibiotic use on the human gut microbiota by comparing individuals who had and had not used antibiotics within the past four months. Wilcoxon tests also were used to compare the average of the shared ASVs and shared genera within individuals and within human and non-human samples in the same households versus other households.

To determine whether sample types significantly differed in microbial community structure, PERMANOVA (Permutational multivariate analysis of variance) tests were used. In addition, we conducted these analyses to identify which variables mainly drive the structure of the human gut microbiota. P values of less than 0.05 (two-sided) were considered statistically significant. To identify genera that significantly differed in relative abundance across sample groups, we used the Analysis of Composition of Microbiomes ANCOM v2.1 R package (50). For declaring differentially abundant taxa, the W-statistic ≥110 were considered as significantly different. The association between antibiotic use and specific microbial taxa was adjusted for the following potential confounders: age, gender and geographical clusters.

## 5. Additional information

## Abbreviations

ASVs: Amplicon sequencing variables,
ARG: Antibiotic resistance genes,
DNA: Deoxyribose Nucleic Acid,
RNA: Ribose Nucleic Acid.

## Funding

This work was supported by Radboudumc Revolving Research Fund (R3Fund) grant and VIDI grant to J.Penders from the Netherlands Organization for Health Research and Development (ZonMw, grant number 016.156.427).

## Author’s contribution

HFLW, JP was responsible for conception, study design, and funding application. All authors contributed to the study protocol development. BVTN led day-to-day management of the study implementation supervised by HFLW, THH, JP and HRvD. HFLW, THH and BVTN took part in getting ethics approval and training for health-care centers. BVTN, MO, and TNNM were responsible for sample collection and other laboratory works. HBH and GG performed the computational analysis. VTVD and BVTN responsible for the statistical data analysis and prepared the figures and tables supervised by JP. BVTN and JP did drafting of manuscript. All authors contributed to the final revision and approved the submission.

## Competing interests

There is no conflicts of interest for all authors.

## Notes

### Competing Interest Statement

The authors have declared no competing interest.

