## Supplemental Figure for "Cross-sectional analysis of the microbiota of human gut and their direct environment (exposome) in a household cohort in northern Vietnam"

**Supplementary.**

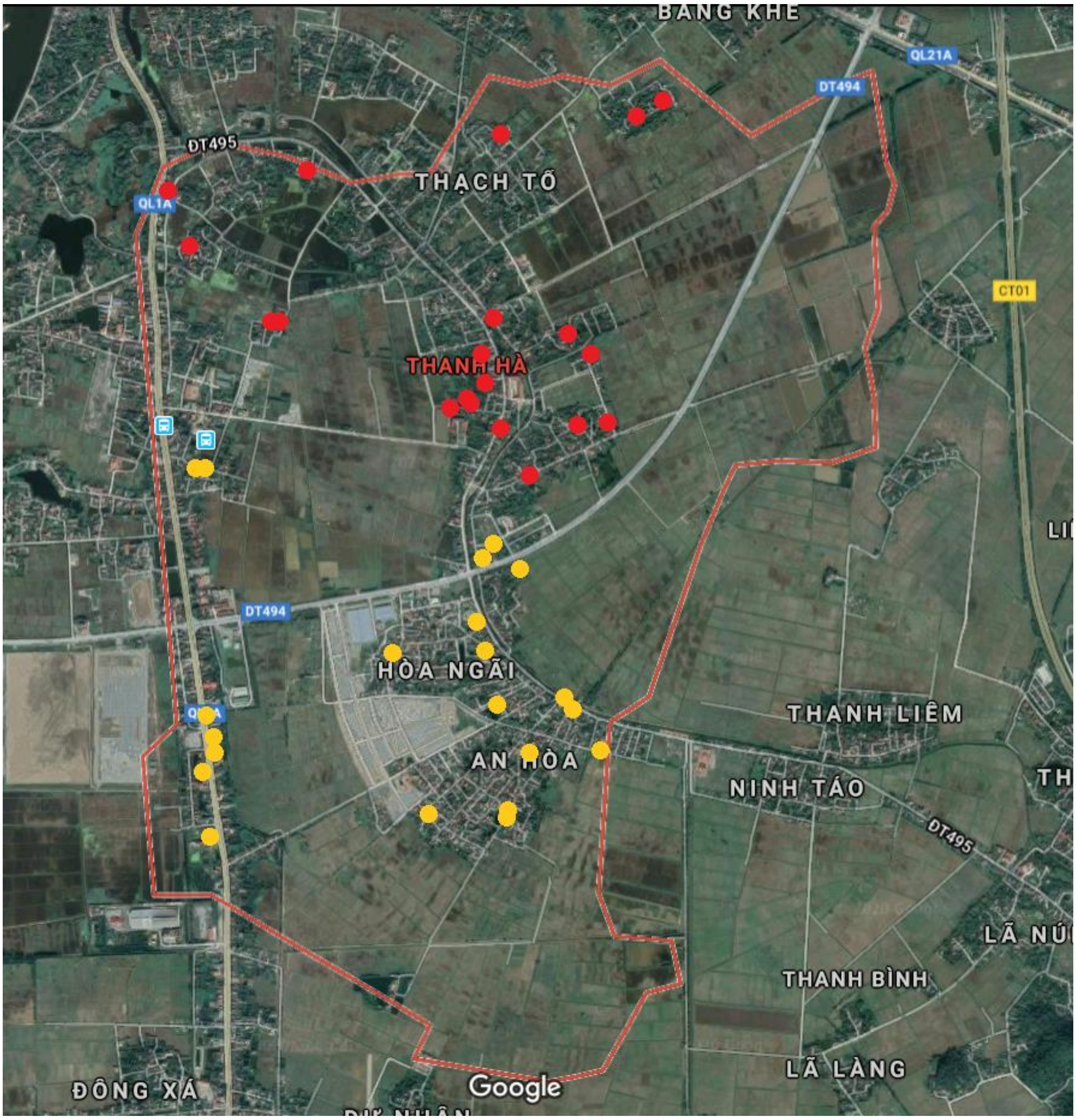


**Figure S1. Geographical location of households.** Each spot represents the location of a household in the community. Red and yellow spots indicate the households, which are belonged to geographical cluster 1 (n=20), and cluster 2 (n=21), respectively. QL1A in blue boxes indicates the 1A national highway; DT494 in blue boxes indicates the 494 provincial road.

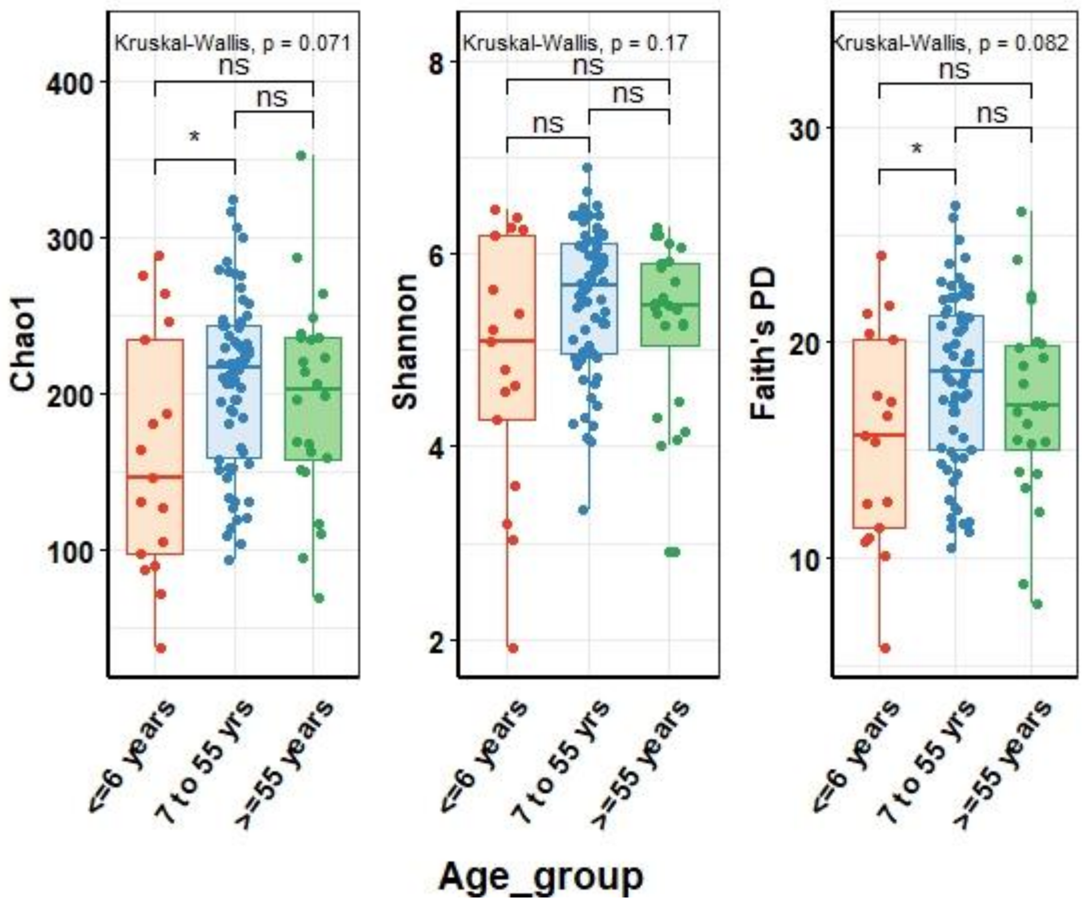


**Figure S2:** Comparison the microbial richness (Chao 1), the bio-diversity (Shannon index) and the microbial diversity that incorporated phylogenetic relationship between ASVs (Faith's PD) in relation to age groups (0-6 years (n=17), 7-54 years (n=79) and ≥55 years and over (n=11)).

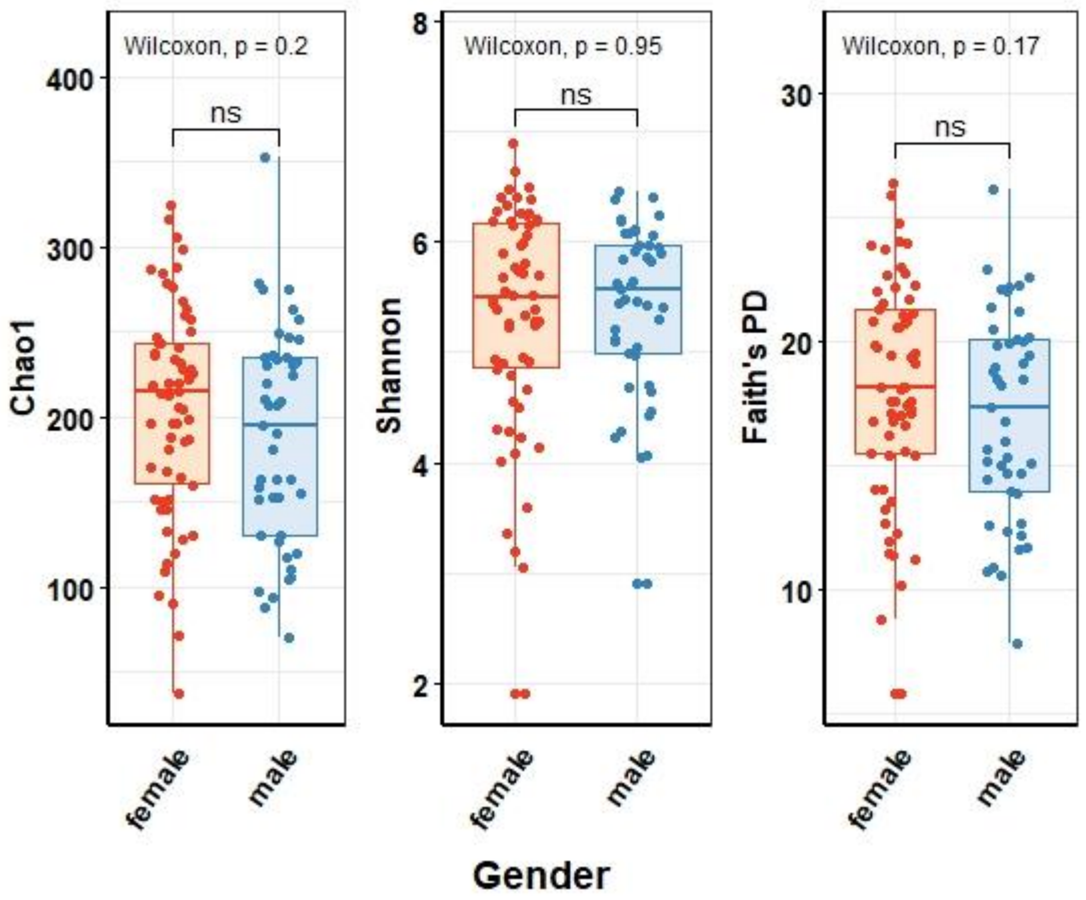


**Figure S3:** Comparison the microbial richness (Chao 1), the bio-diversity (Shannon index) and the microbial diversity that incorporated phylogenetic relationship between ASVs (Faith's PD) in relation to gender of individuals (male (n=45) and female (n=62)).

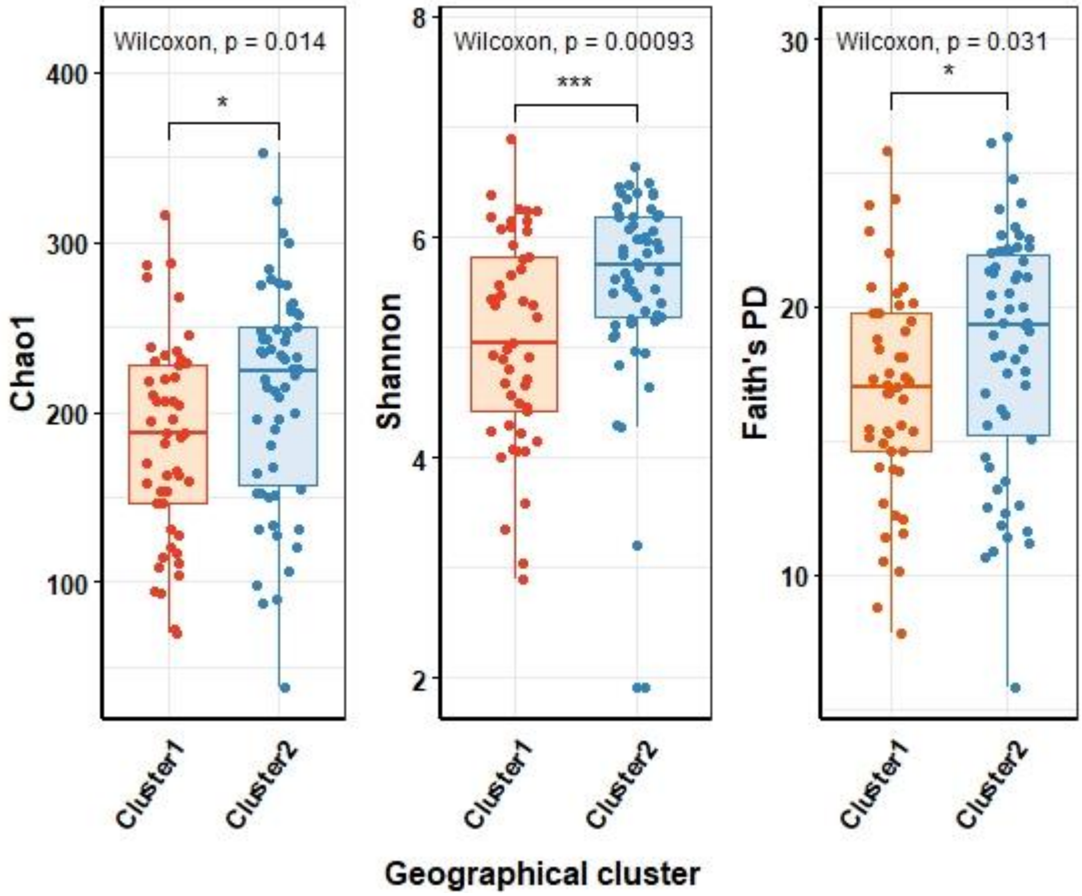


**Figure S4:** Comparison the microbial richness (Chao 1), the bio-diversity (Shannon index) and the microbial diversity that incorporated phylogenetic relationship between ASVs (Faith's PD) in relation to geographical cluster of individuals based on the location of their households (cluster 1 (n=49) and clusters 2 (n=58).

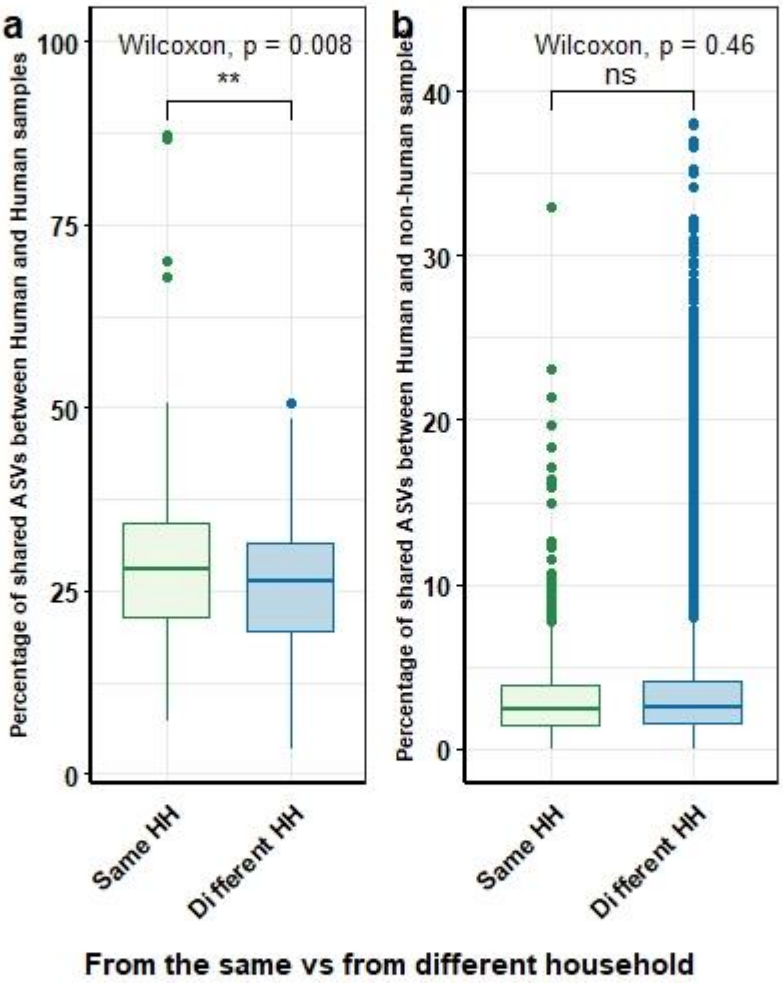


**Figure S5**: Comparison the shared ASVs between humans and between human and non-human samples (Wilcoxon test, ** indicates p-value <0.05, ns indicates p-value >0.05) from the same and from different households (HH),

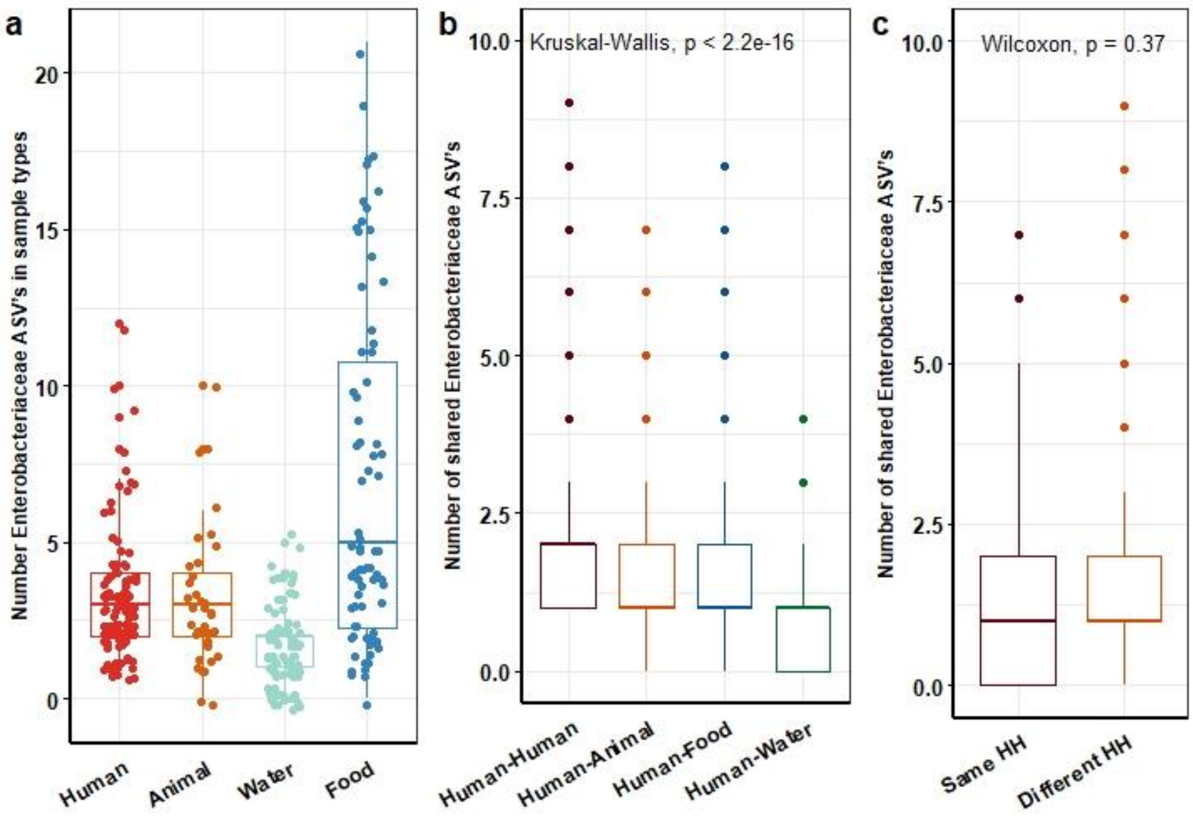


**Figure S6**: Number of ASVs classified as *Enterobacteriacea* in feces from humans (n=107), domestic animals (n=35), water (n=89) and food (n=75) (a). Number of shared ASVs classified as *Enterobacteriacea* between humans and between human versus other sample types (b), comparisons of shared ASVs within the *Enterobacteriacea* genus between human samples and between human and non-human samples from the same and different households (HH) (c).
